# MORTALITY-ASSOCIATED SARS-COV-2 GENOMIC VARIANTS FROM PATIENTS HOSPITALIZED FOR SEVERE PNEUMONIA IN AGUASCALIENTES, MEXICO FROM 2020 TO 2023

**DOI:** 10.1101/2024.03.05.583594

**Authors:** Brian Muñoz Gomez, Miriam Sarahi Lozano Gamboa, Corina Diana Ceapa, Anastacio Palacios Marmolejo

## Abstract

**Background:** The SARS-CoV-2 virus has caused a global health crisis, resulting in a significant loss of human lives. It is essential to report disease and mutation associations to provide ideas for public health interventions and preventive measures.

**Methods:** In this study, to determine the association between genomic variants and the severity of pneumonia caused by SARS-CoV-2, a sequencing analysis of 150 patient samples with confirmed COVID-19 was conducted. These samples were collected between 2021 and 2023 and isolated in Aguascalientes, Mexico.

**Results:** The patient cohort had males and females ranging from 0 to 91 years old. Males accounted for 66% of the population analyzed. The Delta variant was the most prevalent lineage associated with deaths in 2021-2022, while the B.1.1.529 lineages emerged in mid-2022. Currently, the XBB lineage is the most commonly identified in Mexico. New mutations L95M and L46M in ORF 8 and ORF 9 were discovered in 30% and 20% of the sequences and are uniquely present in the studied population. These mutations are positively associated with patient death.

**Conclusions:** This study provides valuable data to aid in understanding the evolution of SARS-CoV-2 in specific populations and explores the severity of the disease and mutation correlations.

## INTRODUCTION

At the end of 2019, a new coronavirus emerged in Wuhan, China. Initially, it was identified as WH-Human 1 coronavirus (WCHV). It exhibited genomic similarity of 79.8% related to SARS-CoV and 59.1% associated with MERS-CoV, as well as showing 100% amino acid similarity in the E and nsp7 with bat SL-CoVZC45 proteins, all possibly derived from recombination events [Wu et al., 2020]. SARS-CoV was determined to have high homologous similarity to CoV bat virus RmYN02 in most of its genome but shows weak homology in the receptor binding domain (RBD) that makes up the S1 subunit of the spicule (S) protein, which represents a critical factor in host receptor recognition and the process of cell membrane fusion with angiotensin-converting enzyme 2 (ACE2). The tissue ubiquity of the latter explains the multiple manifestations and affectations of the disease [Ren et al., 2020].

Coronaviruses, members of the Orthocoronoviridae family within the Coronoviridae domain, are enveloped viruses with a single-stranded positive RNA genome (approximately 26 to 32 kb in length). They are classified into four genera: Alphacoranaviruses (□), Betacoronaviruses (□), Gammacoronaviruses (□), and Deltacoronaviruses (□). The viral genome typically encodes four structural proteins: Spike (s), envelope (E), membrane (M), and nucleocapsid (N), along with numerous non-structural proteins and multiple accessory proteins [Ren et al., 2020].

Now recognized as Severe acute respiratory coronavirus 2 (SARS-CoV-2), it is a highly contagious virus that caused the COVID-19 pandemic disease that brought the world to a standstill, with a primary reproduction number (Ro), which depends on several factors, including the circulating variant, therefore, for Delta the Ro was 3.2 and for Omicron it was 9.5, which also depends on the population density, the mobility patterns, social interactions, and vaccination policies (Roy S, et al, 2023). It is a single-stranded positive RNA virus with a length of 29,891 nucleotides, encoding 9,860 amino acids. Its genome comprises 14 open reading frames (ORFs) and 27 proteins, of which the structural proteins are spike (S), envelope protein (E), membrane protein (M), and nucleocapsid (N) (Chan et al., 2020). As of December 2023, there have been 772 million confirmed cases of SARS-CoV-2 worldwide and 6.9 million deaths attributed to the disease. In Mexico, the first case of COVID-19 was detected on February 27, 2020, and as of today, there are 7.7 million confirmed cases and 334,917 deaths [WHO, 2023; Garcés-Ayala et al., 2020]. The pandemic propelled advancements in molecular biology techniques that allowed us to answer crucial questions, particularly the Delta and Omicron variants. It clarified everything related to the development of the disease, such as the relationship between the characteristics presented and the risk of death or infection susceptibility in prolonged-stay settings [Anderegg et al., 2023; Iqbal et al., 2023].

Epidemiological surveillance has proven indispensable in identifying mutations that affect the viruśs behavior, enhancing its infectious capacity, or evading the immune system.

It has been reported that Latin American countries have experienced an increase in the frequency of virus mutation, with a particular prevalence in these countries compared to other regions. This geographical area could be the origin of new and potentially dangerous virus variants that could later spread globally. This report only covers the period of 2021-2022 [Dhruv Yadav et al. 2022].

In the first year of the pandemic in Mexico, numerous lineages were detected, some transient. In contrast, others, such as B.1, B.1.1.1, and B.1.1.519, among others, represented up to 76% of the circulating lineages in Mexico during this first year, in addition to exhibiting the mutations P314L in protein nsp12 and D614G in spike protein with a prevalence of 99% and 98 % respectively [Taboada et al., 2021]. The spike protein has been a focal point of numeral studies, given its high mutation rate, which gives it the ability to have immune escape. This capability allows the virus to infect even vaccinated or previously infected individuals [Harvey et al., 2021].

After the first year of the pandemic, the Delta variant, with AY20 and AY.26 lineages, replaced other variants and marked the third wave. Finally, the Omicron variant and its respective lineages entered in December 2021, initiating the fourth wave becoming the variant with the highest daily case count. This variant and the recombinant XBB sublineage have spread even among vaccinated populations [García-López et al., 2022]. It is unclear whether the reported regional mutations are related to the alarming case-fatality rates in Mexico. Before vaccination, the rate was 10%, but it has dropped to 4,5% in 2024. These rates are higher than the worldwide average (John Hopkins University, 2024).

Therefore, this project aims to determine the genomic sequence of SARS-Cov2 in samples detected by quantitative reverse transcription polymerase chain reaction (RT-qPCR) in central Mexico from patients with severe respiratory infection to establish a possible association between the severity of pneumonia, death incidence, and genomics variants.

## METHODS

Clinical data were collected from 300 biosamples from deaths in the state of Aguascalientes who tested positive for COVID-19, confirmed by RT-qPCR, spanning from August 2021 to May 2023. The clinical information includes demographic data (age, gender, and vaccination status), comorbidity details, symptoms, and their duration until death (Table 1). Samples with CT values of 28 or less were selected for sequencing using the Illumina CovidSeq protocol following the manufacturer’s instructions and utilizing the Illumina MiniSeq Sequencing System. In total, 173 samples were subjected to sequencing.

**Table 1.** Demographic and clinical characteristics of study subjects.

| n (%) | Total<br>(n=150) | Delta<br>(n=63) | Omicron<br>(n=87) | p-value |
| --- | --- | --- | --- | --- |
| **Demographics** |  |  |  |  |
| Age, years, mean (±SD) | 64 (19.95) | 62 (19.84) | 65(20.00) | 0.068* |
| Male | 99 (66) | 40 (63.4) | 59 (67.8) | 0.581<sup>**</sup> |
| Female | 51 (34) | 23 (36.5) | 28 (32.1) | *** |
| COVID Vaccinated | 41 (27.3) | 14 (22.2) | 27 (31.0) | 0.231<sup>**</sup> |
| **Comorbidities** | 116 (77.33) | 47 (74.6) | 69 (79.31) | 0.496<sup>**</sup> |
| Diabetes | 57 (38) | 21 (33.3) | 36 (41.4) | 0.316<sup>**</sup> |
| Hypertension | 77 (51.3) | 32 (50.8) | 45 (51.7) | 0.910<sup>**</sup> |
| Chronic kidney disease | 18 (12) | 5 (7.9) | 13 (14.9) | 0.192<sup>**</sup> |
| Smoking | 15 (10) | 5 (7.9) | 10 (11.5) | 0.473<sup>**</sup> |
| Heart disease | 7 (4.6) | 2 (3.2) | 5 (5.7) | 0.460<sup>**</sup> |
| Asthma | 3 (2) | 1 (1.6) | 2 (2.3) | 0.758<sup>**</sup> |
| COPD | 8 (5.3) | 4 (6.3) | 4 (4.6) | 0.641<sup>**</sup> |
| Immunosuppression | 10 (6.6) | 0 (0.0) | 10 (11.5) | **0.005<sup>\*\*</sup>** |
| Obesity | 15 (10) | 5 (7.9) | 10 (11.5) | 0.473<sup>**</sup> |
| **Presenting complaints** |  |  |  |  |
| The duration between the appearance of symptoms and death, days mean (±SD) | 16 (9.9) | 18 (11.1) | 14 (8.4) | **0.008<sup>\*</sup>** |
| Fever | 109 (72.6) | 46 (73.0) | 63 (72.4) | 0.934<sup>**</sup> |
| Cough | 115 (76.6) | 54 (85.7) | 61 (70.1) | **0.025<sup>\*\*</sup>** |
| Odynophagia | 63 (42.0) | 27 (42.9) | 36 (41.4) | 0.856<sup>**</sup> |
| Dyspnoea | 135 (90.6) | 58 (92.1) | 77 (88.5) | 0.473<sup>**</sup> |
| Diarrhea | 19 (12.6) | 5 (7.9) | 14 (16.1) | 0.138<sup>**</sup> |
| Chest pain | 33 (22.0) | 17 (27.0) | 16 (18.4) | 0.209<sup>**</sup> |
| Headaches | 96 (64) | 44 (69.8) | 52 (59.8) | 0.204<sup>**</sup> |

\* The p-value is calculated from the Mann - Whitney test, and the \*\* p-value from the chi-square test.\*\*\* p-value is shown in Male cell, Gender was used as a categorical variable.

The obtained FASTQs from the sequencing process were assembled using CLC Genomics Workbench software, with Wuhan-Hu 1 (NC_045512.2) as the reference genome. Subsequently, the Genome Annotation of the BV-BRC platform (https://www.bv-brc.org/app/Annotation) was used to visualize the general genome structure. Based on the quality of each genome, nucleotide contents were analyzed, and those containing 10% or more Ns quantity were excluded from the study. Finally, 150 good-quality sequences were selected for analysis and uploaded to the GISAID platform (https://gisaid.org/) for consultation. Sequence identifiers are stated in Table 2.

**Table 2.**
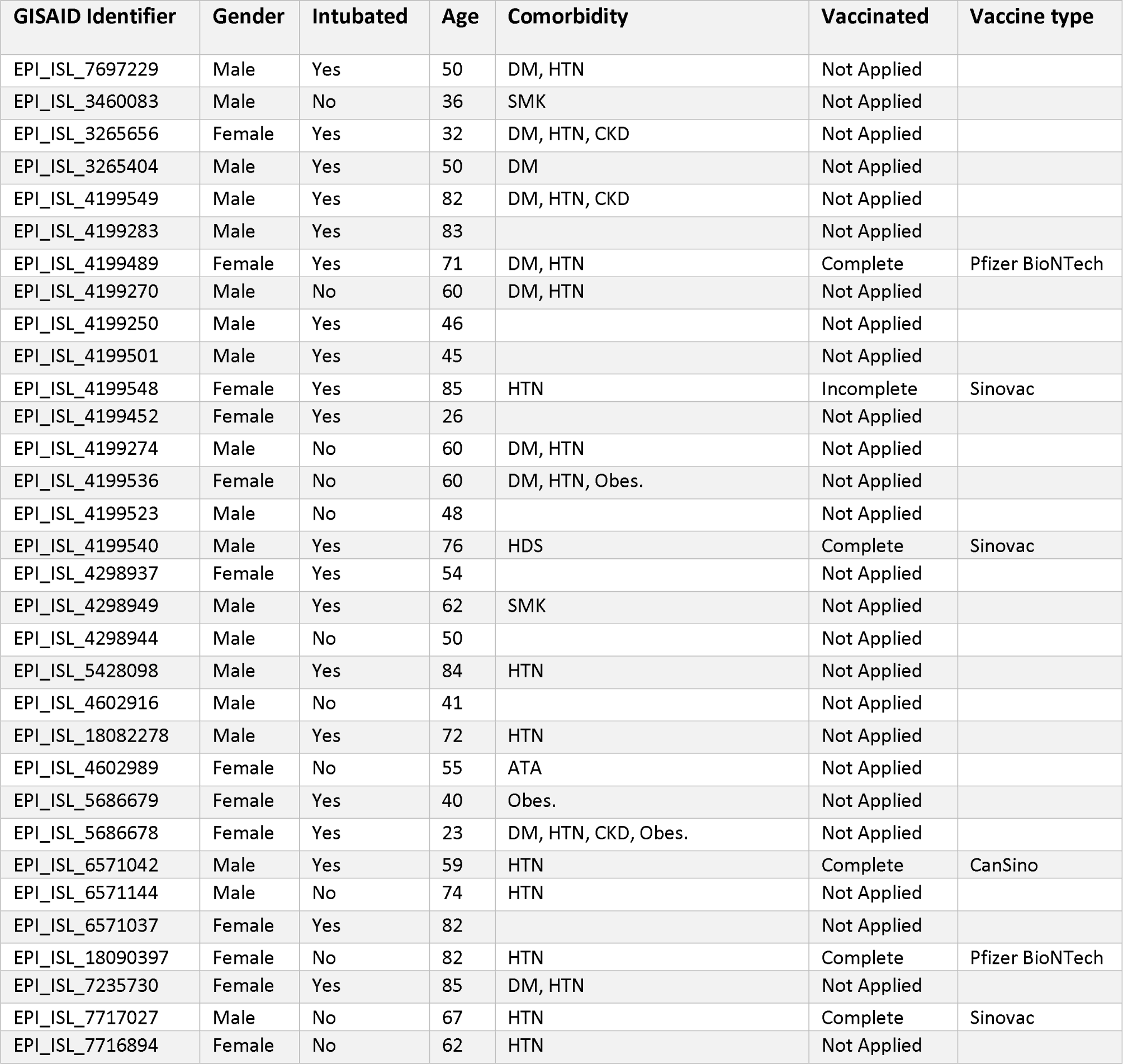
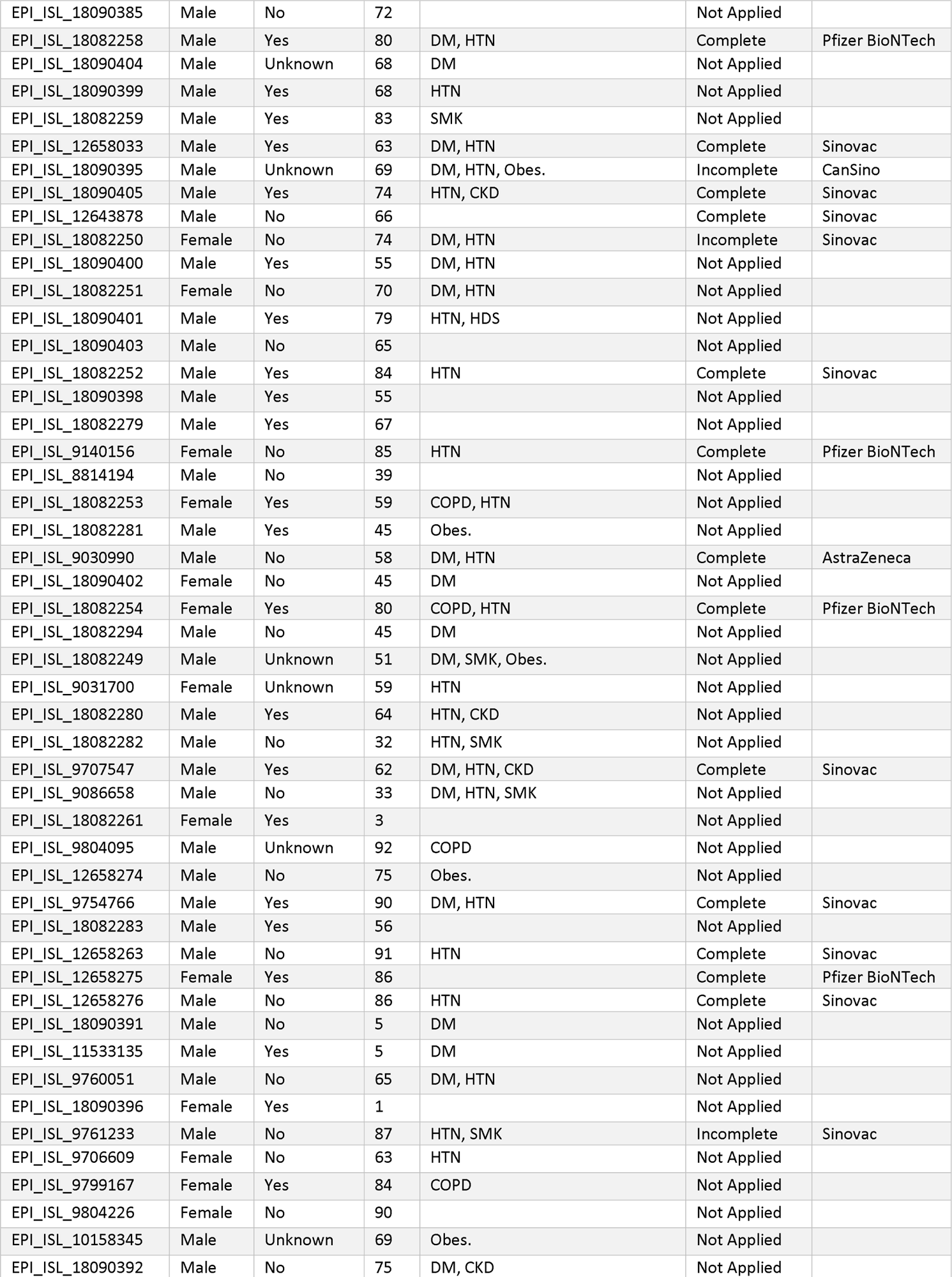
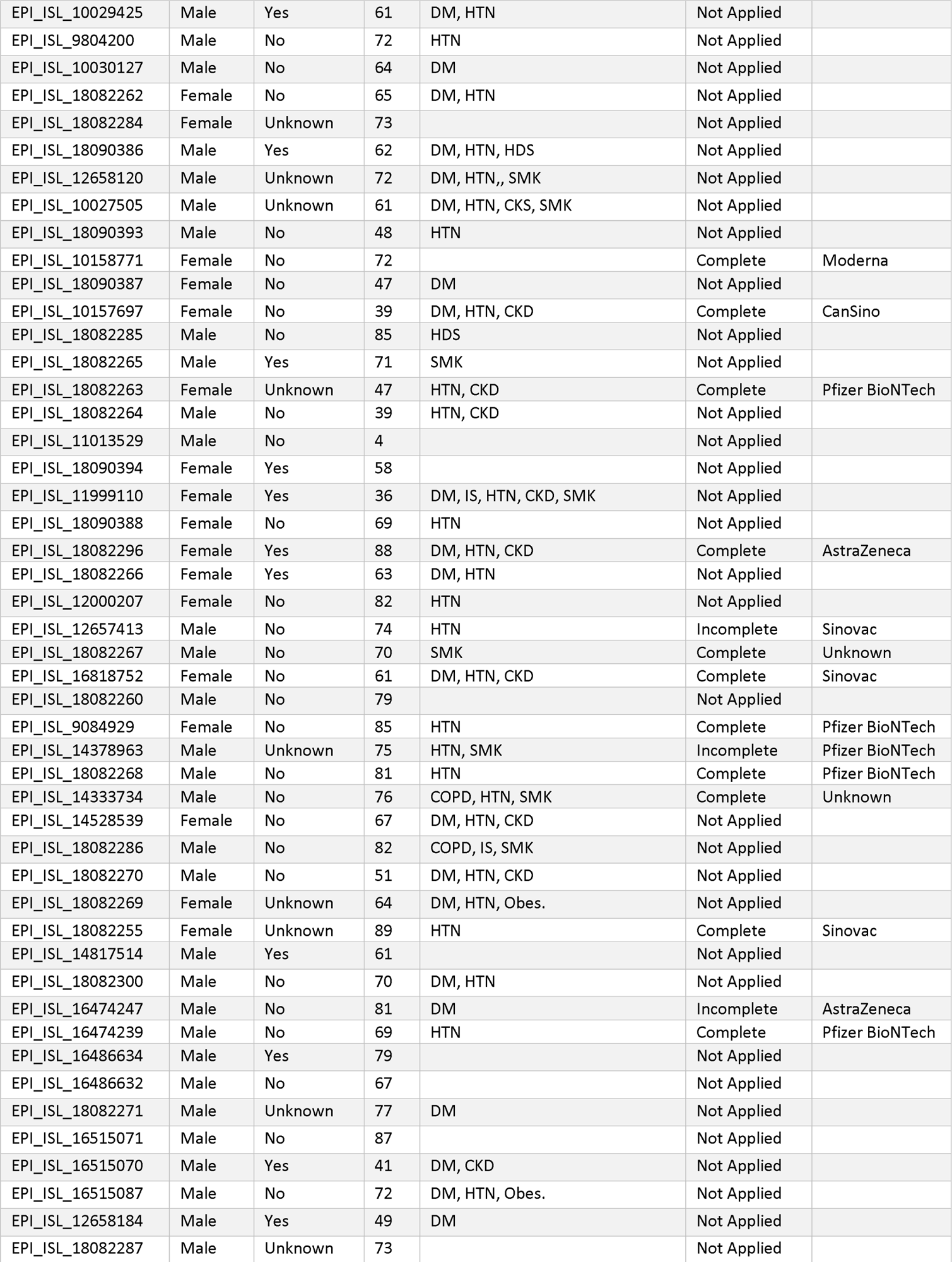
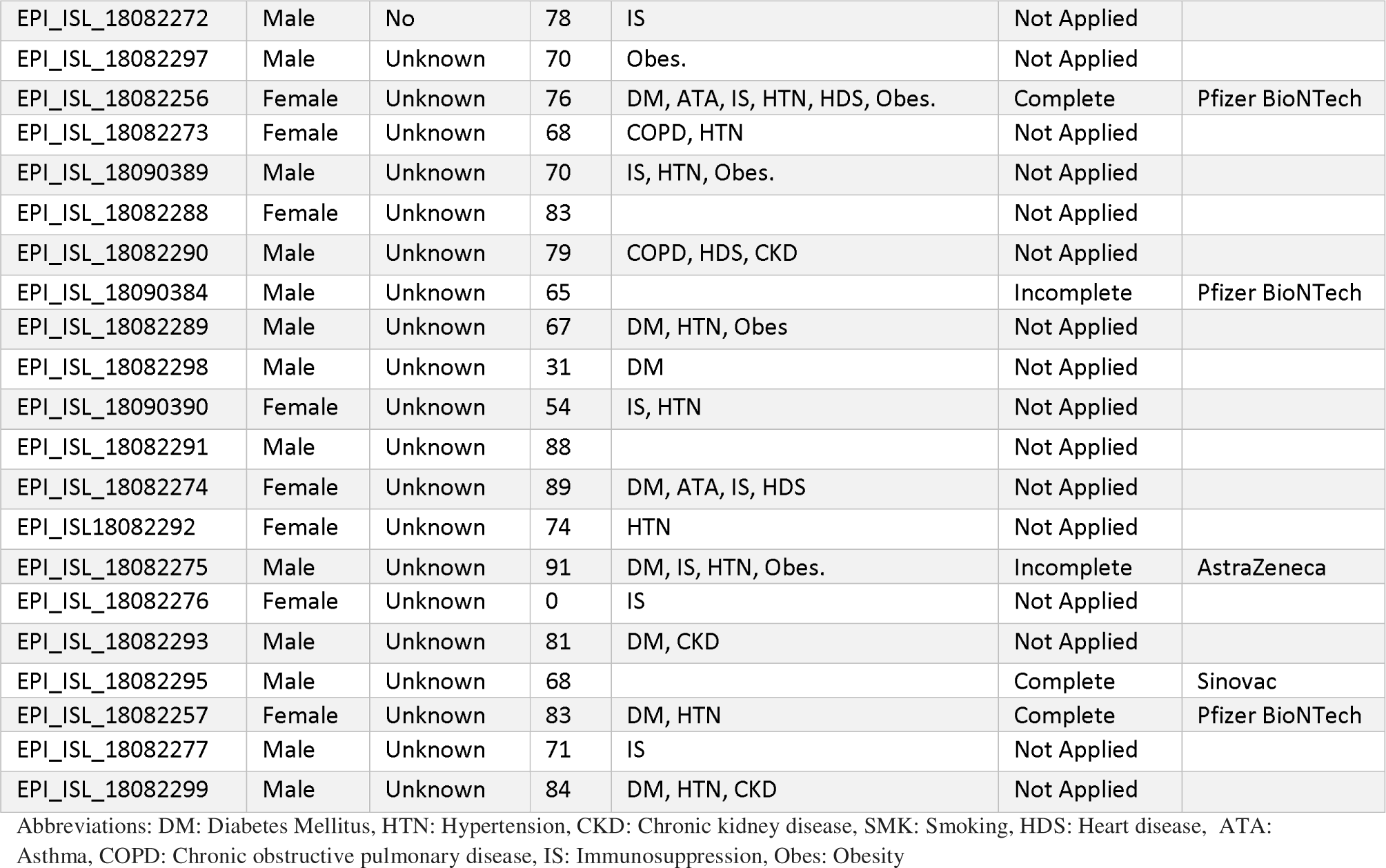
Accession numbers and metadata of viral genome sequenced.

Following the PANGO designation (https://cov-lineages.org/lineage_list.html), the lineage data were interpreted as delta and omicron variants for further analysis. Frequency and percentages were calculated for all the subject characteristics. The mean (±SD) was calculated for the age and duration of the symptoms until death. The normality of data was assessed using the Shapiro-Wilk test. The p-value was calculated through the chi-square test for categorical variables, comparing the presence and absence of each comorbidity and symptom and the male or female gender. Mann - Whitney test was used for non-parametric data comparing the ages of the subjects infected with the delta or omicron variant and the duration of the symptoms until death between the two variants.

Multiple sequence alignments were generated using the online program Multiple Sequence Alignment and SNP / Variation Analysis at the BV-BRC platform (https://www.bv-brc.org/app/MSA) for the different structural and non-structural proteins in different groups and as a whole. The representative genomes of the B, B.1.1.529, BA.1, B.1.351, B.1.617, B.1.1.7, B.1.429, and XBB lineages were used for alignments, using Wuhan-Hu 1 as the reference genome. In the same way, a control group of 443 genomes of the circulating lineages in Aguascalientes between 2021 to 2023 was generated. The genomes selected for the control group are of the period mentioned and at least one sequence of each state of Mexico was introduced.

The alignments were carefully inspected to identify mutations with at least 45% prevalence among the total analyzed sequences. These selected mutations provided an overview of circulating mutations in the center of México between 2021 and 2023. A matrix was made considering the absence and presence of the mutation in each sequence. Using R programming language, a heat map was generated to represent the presence of mutations in each sequence visually. Similarly, the mutations not found in the genome references used in the alignments were considered New Mutations until they were not referenced in official media or previous works. Subsequently, another matrix was used to create a heat map with clustering showing the prevalence of each mutation against lineages from 0 to 100%.

## DATA AVAILABILITY

Sequence data is available on the GISAID platform. All other data is presented in the included tables and figures.

## RESULTS

All patients that were included in this study lost their lives due to the COVID-19 infection or associated conditions. Out of 173 samples analyzed in the genetic analysis, 150 were found to have sufficient quality for further examination and were identified with SARS-CoV2 variants. Of these 150 results, 63 (42%) were identified as delta variants and 87 (58%) as omicron variants. The characteristics, symptoms, and comorbidities of the delta and omicron-infected patients are presented in Table 1. The average age of the patients was 64 years, and there was only a 3-year age difference between averages for the age of the omicron and delta-infected patients. Male patients passed away more frequently than female patients, with 66% and 34%, respectively, and this trend was observed in both groups of variants.

The vaccinated population represented 27%, and further data is needed to determine the effect on disease severity. Hypertension was the most common comorbidity, present in 51% of the subjects, followed by diabetes mellitus at 38%. Asthma was the pre-existing medical condition with the smallest number of cases, occurring only in 3 subjects (2%).

An analysis was conducted to determine the duration between the onset of COVID-19 symptoms and death among infected patients. The mean duration was found to be 16±10 days. However, patients infected with the omicron variant had a shorter mean duration of 14 days, while patients infected with the delta variant had a statistically significant more extended duration of hospitalization by an additional four days (p<0.05). All infected patients exhibited at least two symptoms. Dyspnoea was the most frequent symptom reported by 90.6% of patients. Cough was reported by 115 (76.6%) patients, with 54 and 61 patients infected by delta and omicron variants, respectively, and had statistically significant (p<0.05). Fever was the most frequent complication reported (72.6%), followed by headaches and diarrhea, with diarrhea being the least frequent (12.6%).

Over the three years of study, the evolution of the SARS-CoV2 virus was monitored for epidemiological surveillance in Aguascalientes. Genomic surveillance began in June 2021 and continued until May 2023 (Fig.1a). At the start of the study, the most prevalent lineages in the samples belonged to the delta variant (Table 2), with the AY.20 lineage having the highest number of deaths in the state. However, in early 2022, the prevalence of omicron lineages began to increase, with the B.1.1.529 lineages being the most prevalent. By early 2023, the XBB lineages occupied at least 50% of the COVID-19 cases and rapidly increased to 90% of the sequenced instances over the next few months. Despite this, the BA.1 (belonging to the omicron variant) and AY.3 (belonging to the o delta variant) lineages were responsible for most of the recorded deaths, with 40 and 30 cases, respectively (Fig. 1b).

**Figure 1.**
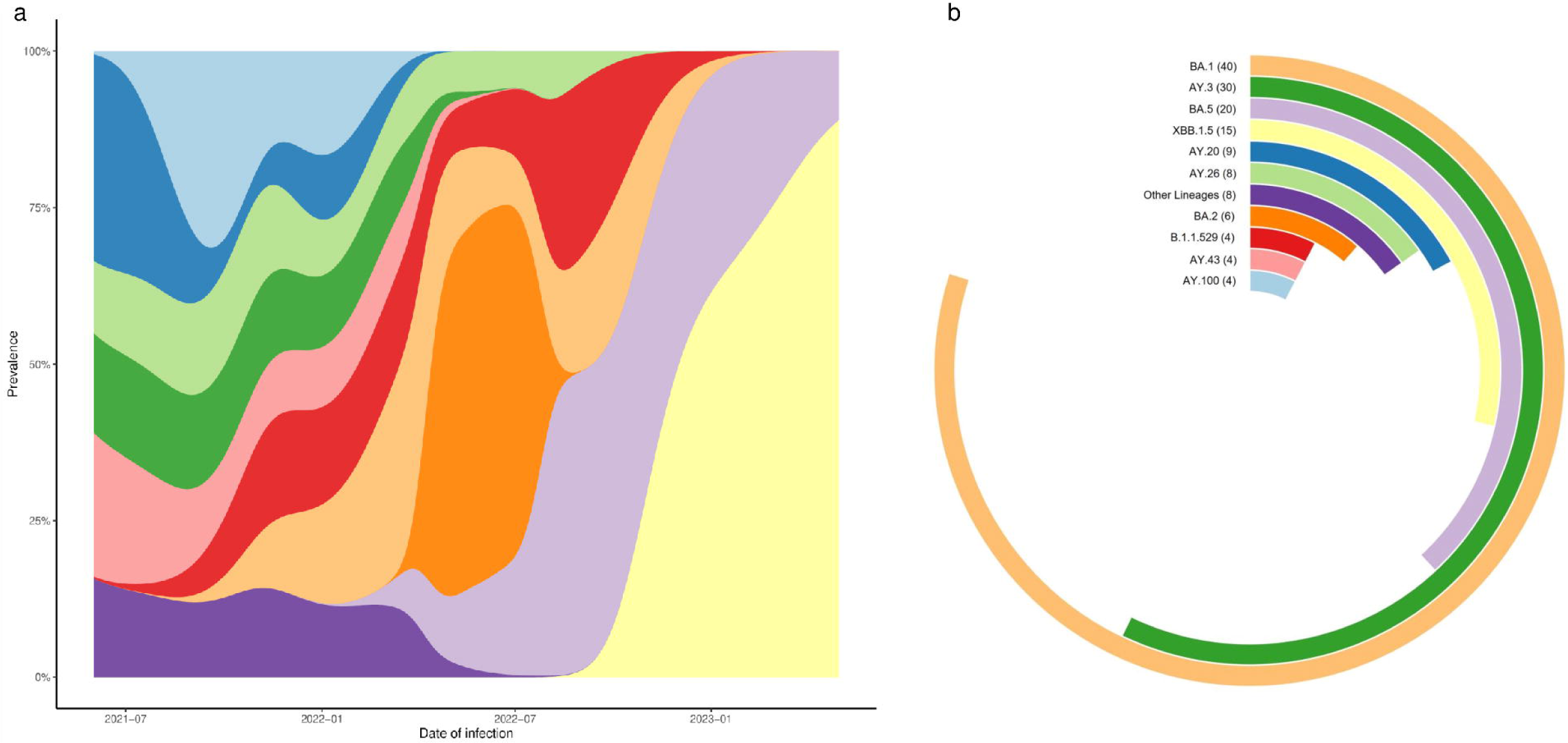
Genomic data in Aguascalientes from June 2021 to May 2023. **a** Lineage prevalence of SARS-CoV2. **b** Number of genomes sequenced from each SARSCoV2 lineage. Lineages for less than one genome are collated into “Other Lineage.”.

The analysis of alignments revealed mutations in all sequences compared to the reference. Mutations present in at least 45% of the sequences were selected for inspection. As expected, sequences identified in 2021 have fewer mutations than those found in 2022 and 2023 (Figure 2). Nevertheless, some sequences belonging to delta variants have more mutations than other sequences of the same lineage (AY.3). Among all the mutations found, the spike protein has the most mutations compared to other structural or non-structural viral proteins. However, the most common mutation in most samples, except for five genomes, is T492I in the nsp4 protein sequence. The mutations P323L for RNA-dependent RNA polymerase (RdRP) and T485K for the spike protein are also present in most sequenced genomes.

**Figure 2.**
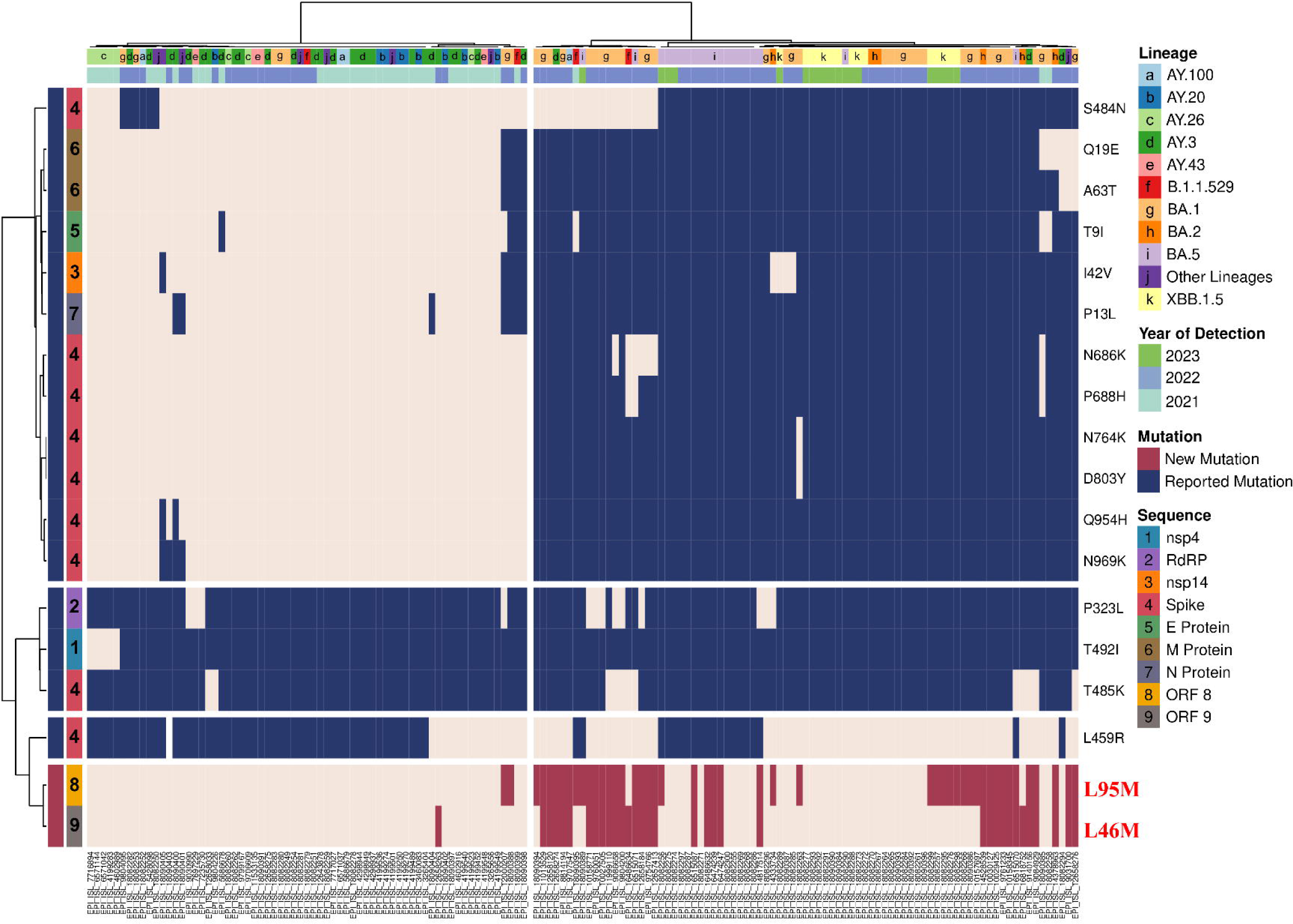
Heat map of the presence and absence of mutations found in each genome sequence of the 150 used. The lineage, year of detection, and mutation sequence have different color codes for better identification. Likewise, the New Mutations show a different color than the previously Reported Mutations.

Additionally, we found two mutations not present in the reference sequences used for different genomes or the control group, shown as New Mutations in Figure 2. The first is L95M for the ORF 8 sequence, representing 31.33%, was found for the first time in the sequence obtained in December 2021, is located in the C-terminal end of the protein (Fig. 5b). The second is L46M for the ORF 9 sequence, representing 22.66%, and appeared in a sample of January 2022, this sample presented both mutations as most of the identified sequences located in the region of interaction with human TOM70 (Fig 5c). Dyspnoea, cough, and hypertension were the clinical data most common in patients whose specimens presented at least one mutation.

The figure 3 shows the absence of these mutations in the control group of 443 sequences from all the states of Mexico, this behavior allows us to assume the discovery of mutations associated with a population. They seem exclusively present in the Aguascalientes state, making this a significant finding for the region. However, further investigations are crucial to determine if these mutations exist in other parts of Mexico. This discovery could have implications for our understanding of genetic variations and their potential impact on human health.

**Figure 3.**
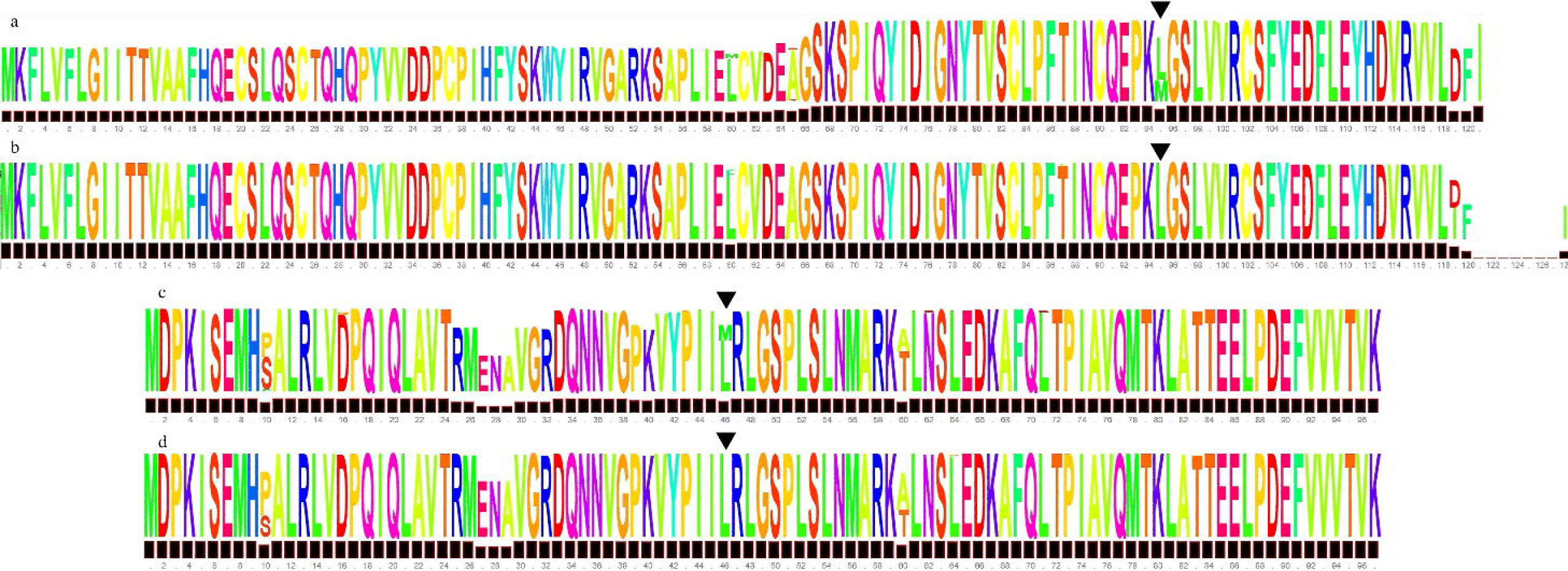
Comparison of LOGO obtained from Multiple sequence alignment (MSA) for ORF 8 and ORF9 sequences between the control and study groups. **a** Schematic LOGO of MSA of the study group highlighting the L95M mutation site for ORF 8 sequence. **b** Schematic LOGO of MSA of the control group highlighting the absence of the L46M mutation for ORF 8 sequence. **c** Schematic LOGO of MSA of the study group highlighting the L46M mutation site for ORF 9 sequence. **d** Schematic LOGO of MSA of the control group highlighting the absence of the L95M mutation for ORF 9 sequence.

As shown in Figure 4, the BA.5 lineage has all the mutations found, showing a prevalence of at least 75% in the corresponding sequences of this lineage. The new mutations have the highest prevalence in the lineages BA.1, BA.2, and BA.5, reaching up to 75% in the corresponding sequences. On the other hand, lineages AY.26, AY.43, and AY.20 have fewer mutations, with only four mutations in all three cases. The mutation T492I for nsp4 is present in all lineages with 100% frequency except for the AY.26 lineage, where it is present in only three genomes, representing 37.5% of the sequences of this lineage.

**Figure 4.**
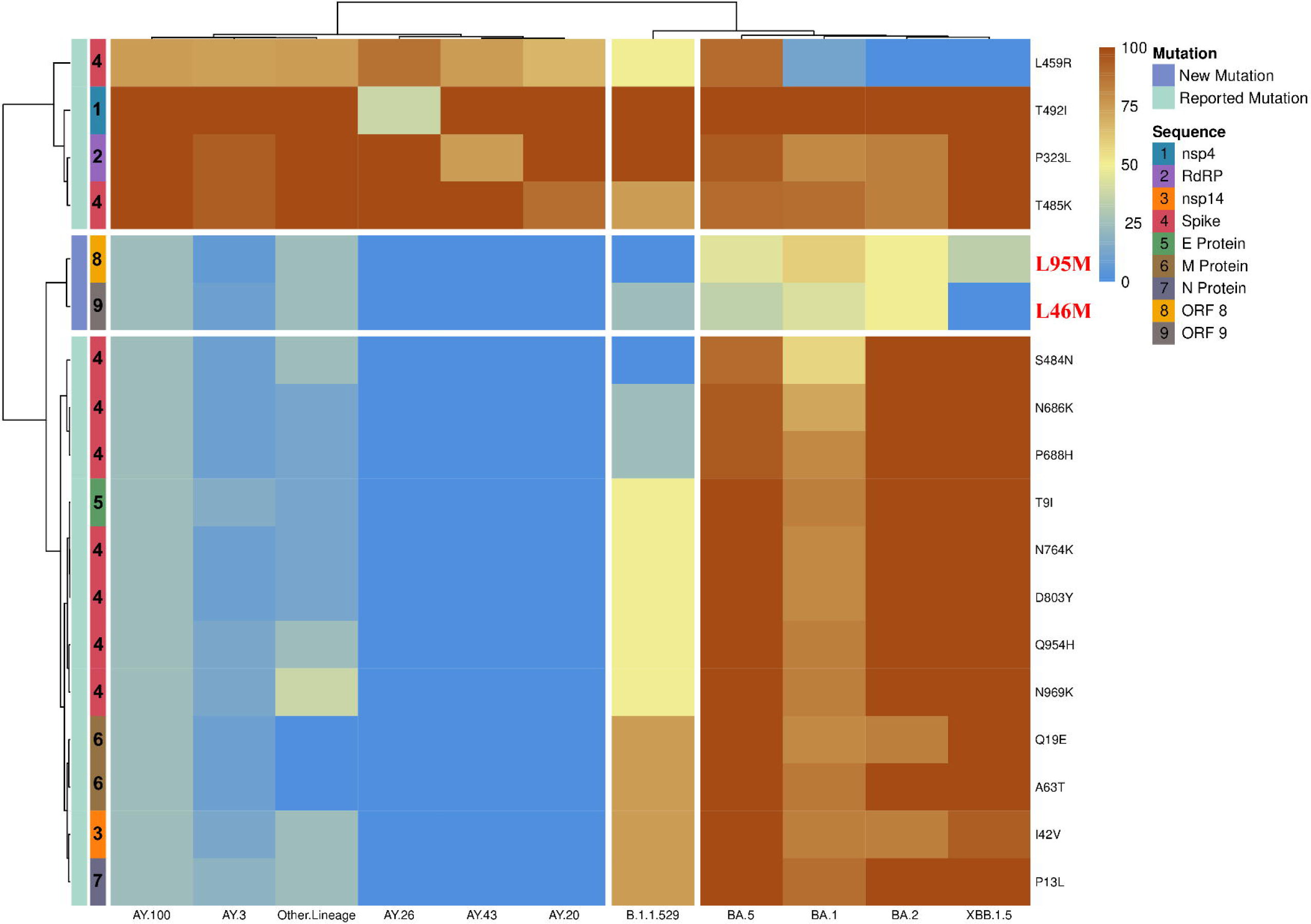
Heat map of mutation prevalence (0 to 100%) in the genomes belonging to lineages found in deaths reported in México between 2021 and 2023. The mutation sequence and the New and Reported Mutations have different color codes for better identification.

The mutation distribution (Figure 5) shows that most mutations are present in the structural and accessory regions of SARS-CoV2. The Spike sequence, in particular, has nine representative mutations out of the 150 genomes analyzed in this study. The graphical genome is a general representation of the SARS-CoV2 genome structure that circulated in the center of México between 2021 and 2023.

**Figure 5.**
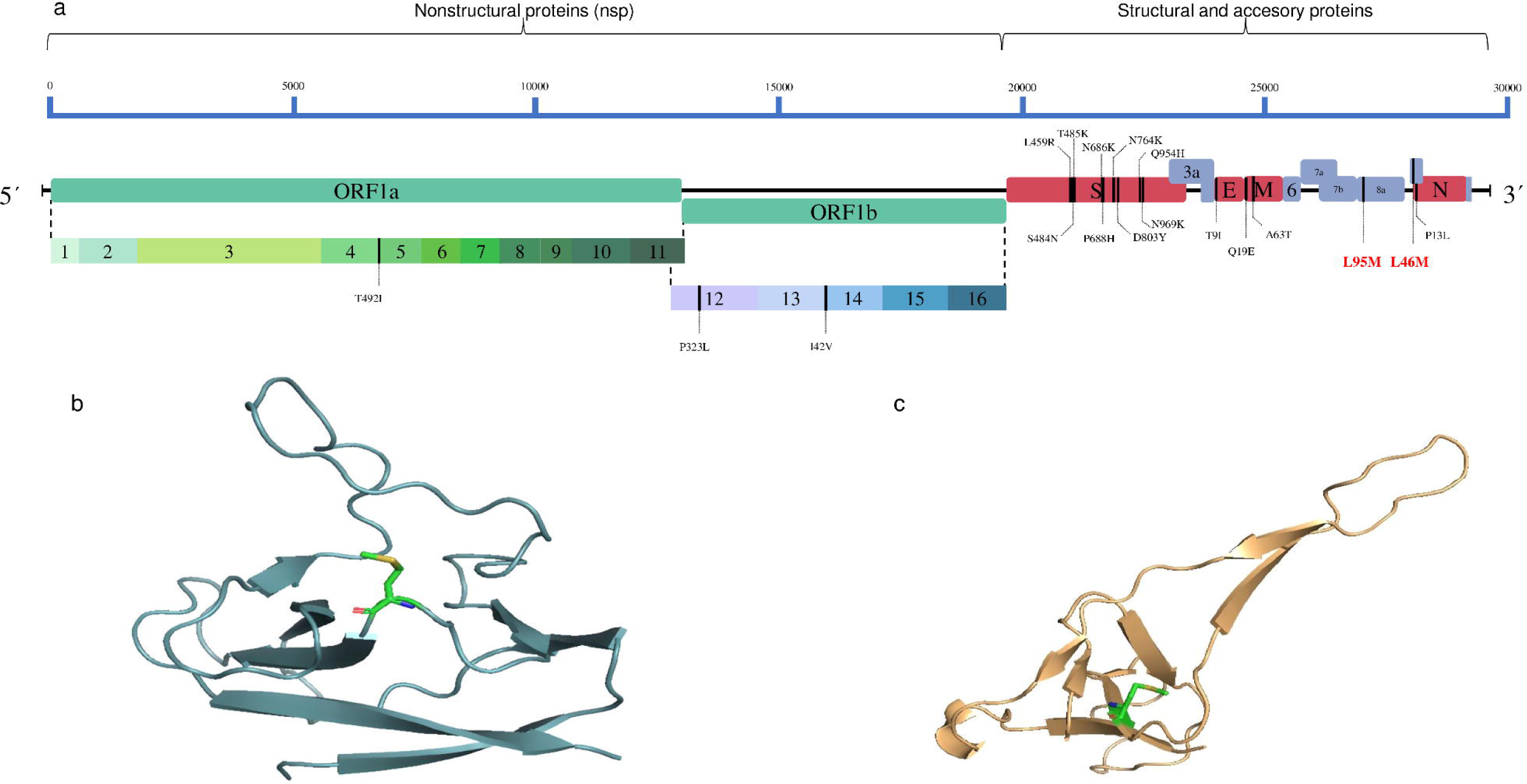
Graphical representation of the SARS-CoV2 and overview of accessory proteins with the new mutations. **a** Genome organization of SARS-CoV-2, the nonstructural proteins are shown in green, the structural proteins are in red, and the accessory proteins are in blue. The mutations are highlighted in their location, with the new ones colored red. **b** Schematic representation of ORF 8 protein with the L95M mutation site highlighted by the amino acid methionine. **c** Schematic representation of ORF 9 protein with the L46M mutation site highlighted by the amino acid methionine.

## DISCUSSION

Mexico established a comprehensive epidemiological follow-up of the SARS-CoV-2 pandemic, carried out by various entities such as public health organizations, academic institutions, and private entities. The collaborative efforts of these entities underscore the significance of a multi-sectoral approach to tackling pandemics of this nature [Hernández-Huerta et al. 2021, Taboada et al. 2021, Taboada et al. 2023]. The World Health Organization (WHO) has confirmed the presence of all Variants of Concern (VOCs) in Mexico as of the latest reporting period. The timely identification of VOCs is paramount for successfully implementing public health measures and mitigating future outbreaks.

According to data from the INEGI (National Institute of Statistics, Geography, and Information), the State of Aguascalientes has an area of 5,680 km^2^, a population of 1.4 million inhabitants, and a population density of 265.2 habs/km^2^, the metropolitan area of the Aguascalientes City concentrates 81% of the total population of the State, people over 60 years of age represent 10.3%. Which undoubtedly represents a risk for the spread of the virus.

According to information from ENSANUT 2018 (National Public Health and Nutrition Survey), abdominal obesity in Aguascalientes reaches up to 72.8% in the adult population, with a prevalence of obesity of 32.6%, high blood pressure of 31.3%, and diabetes of 10.9%. % respectively. In addition, it has an incidence rate of chronic kidney disease of 134.5 per 100,000 inhabitants. (Arreola-Guerra, et al, 2019) being factors that favor the severity of the disease.

The relationship between a host and a virus can be described as co-evolutive, meaning that both entities evolve together over time. This relationship can significantly impact the development of new virus variants, especially in hospitalized patients. This is because when a virus infects a host, it adapts and mutates to survive better and replicate within the host’s body. The host’s immune system also evolves to combat the virus.

However, this co-evolutionary process can become problematic when a virus mutates in a way that allows it to evade the host’s immune system. This can result in the development of new VOCs that are more infectious or virulent than the original virus. This can be particularly dangerous in hospitalized patients, where the immune system may weaken.

Therefore, it is essential to closely monitor the co-evolutionary relationship between hosts and viruses, especially in healthcare settings. By doing so, we can better understand the development of VOCs and take steps to prevent their spread and minimize their impact on public health.

In Mexico, from 2020 to March 2022, 10% of COVID-19 patients required hospitalization, and the lethality rate was less than 1% [García-López et al. 2022]. Hospitalized patients had a higher lethality rate of around 40%. Among these hospitalized patients, around 1% required mechanical ventilation and had an even higher lethality rate of around 80%. Analyzing patient demographics, age, and patient type are crucial factors in COVID-19 outcomes in Mexico [Torres-Ibarra et al., 2022]. Patients under the age of 65 who require hospitalization have a mortality rate of less than 50%, which increases significantly with advancing age. Patients aged 85 years or older have the highest mortality rate, reaching up to 60% [García-López et al., 2022]. Connecting specific virus mutations to particular cases is challenging due to their rarity.

However, the State Public Health Microbiology Laboratory (LESP) in Aguascalientes has the advantage of having access to such valuable samples and the ability to monitor the corresponding mutations in this particular population continuously. This unique advantage allows for a comprehensive analysis of the genetic variations of the virus. This analysis can significantly contribute to understanding its regional evolution and spread.

This work focuses on molecular epidemiology and COVID-19-infected patient demographics for all pneumonia-associated deaths in the Aguascalientes state from the pandemic’s beginning until May 2023. Aguascalientes is a small central state situated in the Mexican Bajio region, serving as a connecting hub between various large states in the center of the country, including San Luis Potosi, Queretaro, Guanajuato, Zacatecas, and Nayarit due to its leading económic sector being the manufacturing industry. The state is located in the Mexican Plateau, which connects Mexico’s Central and North regions. Previous country-wide molecular epidemiology efforts have included this region, but only a few sequences were analyzed compared to other states, such as the capital.

The CDC has reported that severe COVID-19 infection is more likely to occur in patients with underlying chronic diseases, with 90% of hospitalized COVID-19 patients having at least one chronic condition. Hispanics, who have a higher burden of chronic diseases such as obesity, diabetes, and renal disease, are at greater risk for severe COVID-19 outcomes [Gil et al. 2020]. Worldwide, comorbidities such as obesity and diabetes may be associated with a younger age of severe disease in the Mexican population [Herrera-Esposito et al. 2022]. In accordance, in the study group, the most prevalent co-morbidities were diabetes and hypertension.

It is essential to consider that mortality rates are low in younger people, which we also see here. The average age of the studied group is 64 years, significantly higher than the publicly reported average age of 44 years in the Mexican population. This is because older age is associated with worse health outcomes and higher death rates for COVID-19 patients.

On average, in Mexico, it was previously reported that the delta variant infection led to the highest maximum viral load and shortest time from symptom onset to maximum viral load [Ribeiro et al., 2023]. There were similarities in the genomic sequences of SARS-CoV-2 found in Mexico, Belize, and Guatemala in 2021, which are associated with a highly infectious and virulent strain [Hernández-Huerta et al. 2021]. This demonstrates the rapid virus mutation rate during the pandemic’s first months to one year. Our results corroborate these findings and show that the mutation rate has not slowed from 2021 to the end of 2023.

This study acknowledges that the aforementioned general aspects of the infection apply to our data. The patient population had an average age of 64 years. The mean age of patients infected with the Omicron variant was only three years different from those infected with the Delta variant. Male patients exhibited a higher mortality rate than female patients, with males accounting for two-thirds of the analyzed population as compared to females.

This study could also confirm that children who are immunocompromised and under the age of three who have contracted COVID-19 and have comorbidities, particularly diabetes or obesity, are at a greater risk of developing pneumonia, leading to death.

This study also confirms previous findings that BA.5 Omicron variant dominated the fifth epidemic wave (summer 2022) in Mexico, replacing BA.1 and BA.2 and that Aguascalientes was among the states with a transitory presence of BA.2 and a low presence of BA.4 [Taboada et al. 2023, Castelán-Sánchez et al. 2023, [García-López et al., 2023].

The most intriguing discovery involves the identification of two mutations that appear to be associated with the population of Aguascalientes. Both proteins are linked to immune system evasion [Zandi et al. 2022]. The first, L95M in ORF8, has demonstrated a high mutation rate at amino acids 119 and 120, conferring resistance to evasion. Nevertheless, this mutation is located in the region of highest diversity found in North America and Oceania, despite not being previously reported [Alkhansa et al. 2021]. On the other hand, ORF9 protein has shown interaction with TOM70 (mitochondrial import receptor), reducing active IFN. The L46M mutation is within the 11 amino acid residues that form the binding complex with TOM70 [Gao et al. 2021], making it necessary to investigate whether it has a more significant impact on this binding and, therefore, a more significant role in the immune response. These findings open the door to analyzing the effects of these mutations and their potential implications for viral infectivity, disease severity, or clinical characteristics.

## CONCLUSIONS

After evaluating patient demographics, virus evolution dynamics, and changes over time, we identified unique characteristics of SARS-CoV-2 infection in patients hospitalized for severe pneumonia in central Mexico during the pandemic. Our findings revealed two new mutations specific to the lineages in this geographical area. We also observed only two prevalent SARS-CoV-2 genomic lineages in 2023. Continuing genomic surveillance is crucial to detect emerging variants that could potentially threaten public health. This study emphasizes the significance of maintaining such vigilance to safeguard our communities. Understanding the virus’s genetic makeup and evolution patterns is essential for developing effective vaccines and treatments.

## AUTHOR STATEMENTS

### CONTRIBUTIONS

BMG, MSLG, CDC, and APM developed the idea and wrote and edited the manuscript. BMG MSLG performed laboratory and bioinformatics analyses. BMG prepared the manuscript figures. We acknowledge using Grammarly AI to improve the manuscript’s English writing [Grammarly 2024].

### CONFLICTS OF INTEREST

The authors declare no conflicts of interest. All authors have submitted the ICMJE Form for Disclosure of Potential Conflicts of Interest.

### FUNDING INFORMATION

The authors declare that no funding was received for this project.

## Notes

### Competing Interest Statement

The authors have declared no competing interest.

